# *Dlic1* deletion impaired cerebellar development in mouse

**DOI:** 10.1101/176339

**Authors:** Shanshan Kong, Wufan Tao

## Abstract

The cerebellum is an important model system to study Central Nervous System, it has a striking morphology consisting of folia separated by fissures of different lengths, and the molecules controlling the process of cerebellar development have been unclear. In this study, we report an unrecognized contribution of *Dlic1* to the postnatal development of murine cerebellum. *Dlic1* encodes a light intermediate chain of cytoplasmic dynein 1, it is a subunit unique to the cytoplasmic form of dynein, and how it contributes to dynein function is not fully understood. Our data show that *Dlic1* deletion leads to impaired cerebellar development, foliation and fissuration, the volume of the cerebellum was evidently reduced. Consistent with the histological defects, *Dlic1* knockout mice show defective motor coordination, this strongly indicates the fundamental role of *Dlic1* in cerebellar development and function. Purkinje cells are located within the cerebellum and form synapses with granule cells. In *Dlic1*^*-/-*^ mice, Purkinje cells display reduced dendritic complexity and the distribution of dynein in Purkinje cell layer is changed. We also found that *Dlic1* deletion reduced the protein levels of dynein subunits and leads to the defective assembling of dynein subunits. These data indicate that *Dlic1* may affect cerebellar development through affecting the normal function of dynein.

**Abbreviations:**

*Dlic1*: cytoplasmic dynein 1 light intermediate chain
CNS: central nervous system
MEF: mouse embryo fibroblast

## 1. Introduction

The cerebellum is an important coordination center that controls the motor activity in our brains, its volume just represents 10% of the total brain volume, but its neuron numbers account for half of the total neurons in the brain[1]. The cerebellar growth and pattern formation is one of the most dynamic events during brain morphogenesis, and the developmental defects in cerebellum can lead to serious human disease. But people know surprisingly little about cerebellar development. The cerebellar structure divided into three longitudinal regions: the vermis, the paravermis and the hemisphere[2]. Each of the regions is folded into a lube. The malformation of lubes will lead to the human congenital ataxia, which is appeared in early life[3]. Cerebellar development is an exquisitely orchestrated process that produces a layered structure of the cerebellar cortex containing granule cells, Purkinje cells, and cerebellar interneurons. During cerebellar development, Purkinje cells develop extensive dendritic arbors and synapse onto granule neurons and regulate the proliferation of granule cells[2], [4]. The crosstalk between granule cells and Purkinje cells is important for driving the proliferation of granule cells and formation of the mature architecture in the developing mouse cerebellum[5].

The axonal transport of organelles is critical for the development, maintenance and survival of neurons, and its dysfunction has been implicated in several neurodegenerative diseases. Retrograde axon transport is mediated by the motor protein dynein. Neurons including Purkinje cells and granule cells are highly polarized cells composed of dendrites and axons[6]. During the development of axons and dendrites, the localization of intracellular signaling proteins to specific compartments is acquired[7], since the localization of signaling proteins to distinct subcellular regions modulate signaling outputs[8]. Microtubule-based axonal transport is critical for the development, maintenance and survival of neurons. During development, the axonal transport of Golgi-derived vesicles, synaptic vesicles, mRNA granules, mitochondria and other organelles is considered necessary for appropriate axon extension, guidance and branching.

Cytoplasmic dynein, the microtubule minus end-directed motor protein, is essential for many cellular functions(Hirokawa et al., 2010). In neurons, dynein is responsible for the retrograde transport of cargos, such as signaling endosomes, from the axon terminus to the cell body. Retrograde transport is required for neuronal survival and axonal plasticity(Van Aelst and Cline, 2004). And mutation in dynein can lead to numerous neuron degenerative diseases [11]–[14]. The function and physiological importance of dynein in the cerebellum have not been defined. Since dynein has two different light intermediate chains: DLIC1 and DLIC2, the distinct functions between the two pools of dynein still remain unclear.

Our previous work has found that *Dlic1* is highly expressed in central nervous system[15], the global deletion of *Dlic1* in mouse appeared normal at birth and displayed retinal degeneration. In this story, we found that the *Dlic1* is required for the neuron development in the cerebellum. The development of the granule cells and Purkinje cells are impaired in *Dlic1^-/-^* cerebellum, and *Dlic1* deletion affected the subunit protein level and then localization of dynein in neurons.

## 2. Materials and methods

### 2.1 Animals

All the *Dlic1* mutant mice used for experiments were described by S. Kong et al[16] and maintained on a C57BL/6 and 129/sv mixed genetic background. During the experiments, the heterozygote *Dlic1^+/-^* littermates were used and processed in parallel with the *Dlic1^-/-^* mice for comparison purposes. All animal experiment procedures were performed in accordance with the protocols approved by the Animal Care and Use Committee of the Institute of Developmental Biology and Molecular Medicine at Fudan University.

### 2.2 Neuronal Culture

Granule cells were dissociated from P3 mice after birth. After trypsin digestion, dissociated cells were plated on cover glasses coated with poly-D-Lysine (Sigma) and were cultured in Neuro basal Medium (Invitrogen) supplemented with 2% B27 (Invitrogen), 10% FBS, 1% penicillin/streptomycin, and 1% L-Glutamine[17]. After 36–48hrs in culture, cells were fixed in 0.4% PFA at room temp for 10 min and subjected to immunostaining. The mean values were calculated from three separate experiments.

### 2.3 Behavioral Analysis

Footprint analysis was carried out as described previously[18]. Briefly, 3 month old mice with hind limbs coated ink were allowed to walk on papers through a tunnel (50 cm long, 9 cm wide, and 16 cm high). The subsequent footprint patterns were evaluated by step length and gait width; step length is the average longitudinal distance between alternate steps, gait width is the average lateral distance between left and right steps measured by the perpendicular distance of a given step to a line connecting its opposite preceding and succeeding steps.

### 2.4 Histology and Immunostaining

Postnatal brains were dissected and fixed by immersion in 4% (w/v) paraformaldehyde overnight at 4°C. Tissues were embedded in OCT according to standard methods and frozen sectioned at 10μm. For histological analysis, sections were stained with hematoxylin solution for 5 min, counterstained with 0.5% (v/v) eosin alcohol solution for 2 min, dehydrated, and mounted. To quantify cerebellar area, digital images were taken under light microscopy, five sections in the central and midline were counted for each cerebellum. Three mice for each time-point and each genotype were used for statistics.

For Immunostaining, frozen sections were stained with various antibodies as standard protocol. The primary antibodies used were: mouse anti-calbindin (C9848; Sigma), DIC (74kDa) (MAb1618, Millipore); mouse anti- EB1(010811B11, Absea) Secondary antibodies: goat anti-rabbit IgG-FITC (F9887) and goat anti-rabbit IgG-CY3 (C2306) were from Sigma; goat anti-mouse IgG-FITC (FI-2020) was from Vector Labs. All the secondary antibodies were diluted at 1:300 when used.

Purkinje morphology was assessed by dendritic parameters using a modification of the method described by Sholl [19]. To calculate the dendrites, 5 fields of Purkinje cells in the cerebellum (lobules V – VIII) were randomly imaged by confocal microscopy (Zeiss Axiovert 100 M). Total dendrites were traced and analyzed by the Image J software with Neuron J plugin.

### 2.5 Sucrose gradient centrifugation

Total mouse brain protein were extracted with lysis buffer (10% glycerol, 50 mM Tris–HCl, pH 7.4, 10 mM MgCl2, 2 mM ATP, 0.5 mM DTT, 100 mM NaCl and protease inhibitors) through French press. The whole brain lysate was loaded onto a 5–20% sucrose gradient prepared in lysis buffer. The tubes were ultracentrifuged at 304,660g for 16 h at 4 °C. 1mL fractions were collected, and proteins were resolved on SDS–PAGE and blotted with anti-DIC (74 kDa) (MAb1618, Millipore) and anti-EB1 (010811B11, Absea).

## 3. Results

### 3.1 Dlic1 deficiency in mice impaired the cerebellar development

*Dlic1* null mice show growth retardation and abnormalities in retina[16]. To clarify the role of *Dlic1* in cerebellar development, we examined the morphology of cerebellum in *Dlic1^-/-^* mice (Figure1). Microscopic examination showed the cerebellar size is reduced at both P3 and P14 when *Dlic1* is deleted (Figure1 A, B), the anterior-posterior axis of cerebellum are shorter compared with the control mice. Additionally, the *Dlic1^-/-^* mice also experienced overall brain weight loss at P3 and P14 (Figure1C). The reduced anterior-posterior axis (AP) length of the *Dlic1^-/-^* cerebellar central region (Figure1 A, B) suggesting defects in cerebellum foliation. To closely examine the foliation of cerebellum in *Dlic1^-/-^* mice, we analyzed the cerebellar morphology and the cerebellar midline area in H&E stained frozen sections (Figure2). From the section analysis, we can see the overall cerebellar size and foliation are severely affected in *Dlic1^-/-^* mice in a series of stages: at P1, a mild size reduction, and foliation defects can be detected (Figure2A); at P7 and P14, the cerebellar size is continued to reduce and the foliation defects become more evident (Figure2B, C). The lobules VI and VII are severely hypoplasia and fused together in *Dlic1^-/-^* cerebellum (Figure2B, C). The total number of lobules is reduced when dlic1 is absent. These results show that *Dlic1* plays an important role in cerebellar development, deletion of *Dlic1* leads to reduced cerebellar size and defective foliation.

**Figure1.**
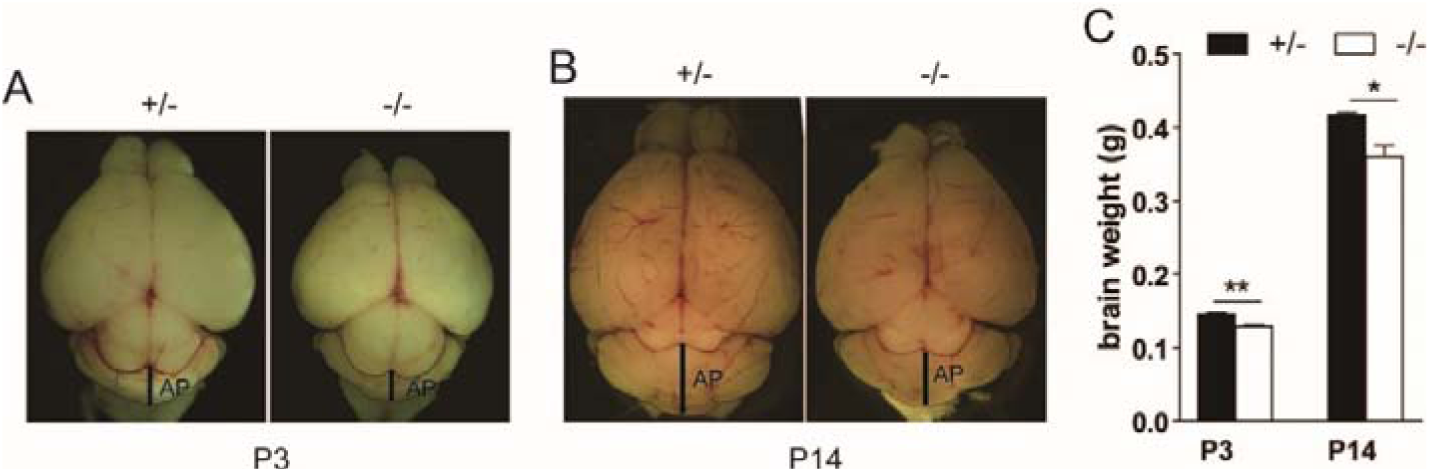
Reduced cerebellum size in *dlic1^-/-^* mice. (A-B) Whole-mount images of representative mutant and littermate control brains at P3 and P14. The cerebellum were outlined with black lines and the anterial-posterial axis were labeled with AP. (C) Gross analysis of the mutant and littermate control brains at P3 and P14. (n=3 for each genotype,*, P<0.05, **, P<0.001)

**Figure2.**
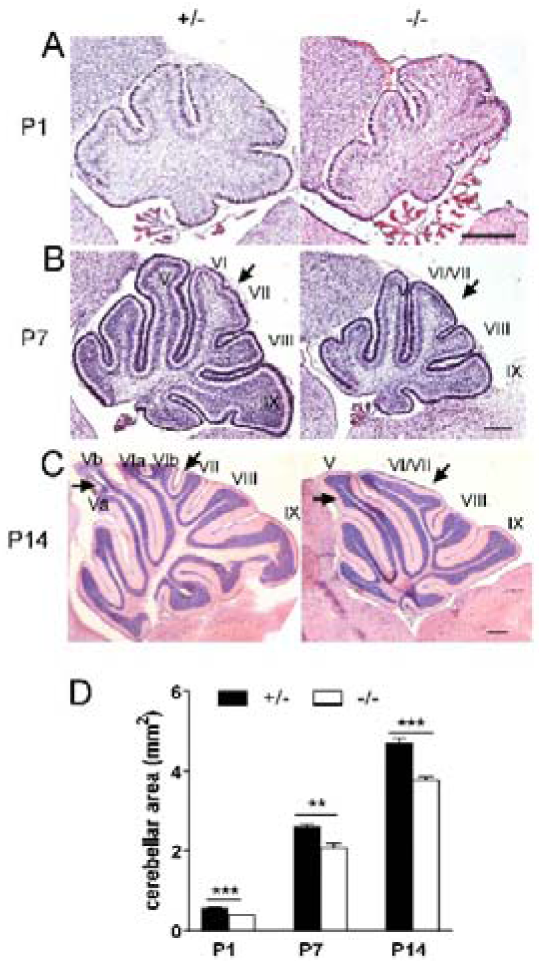
Cerebellar development defects in *dlic1^-/-^* mice. (A-C) Midline sagittal sections of cerebellum from representative littermates at P1 (A), P7 (B) and P14 (C). Folia are labeled, and arrows indicate foliation defects: decreased size of V and fusion of VI and VII. Scale bar: 200 μm. (D) Quantification of cerebellum area of midline sections. **P < 0.01, ***P < 0.001, n = 3 for each genotype

### 3.2 Dlic1^-/-^ mice display impaired motor coordination

The histological differences described above prompted us to investigate whether the *Dlic1* is involved in cerebellar motor function. Footprint analysis is a well-established and widely used protocols for measuring motor coordination and balance in mice and rats. Three age-matched *Dlic1^+/-^* and *Dlic1^-/-^* were coated the hind feet with nontoxic black ink and walk on paper along a 50-cm-long, 9-cm-wide runway, with 16-cm-highwalls on either side. Seven consecutive steps were recorded in terms of step length (Figure3B) and width (Figure3C). Foot print analysis shows that *Dlic1^-/-^* mice exhibit reduced gait length and the gait width at 3 months compared with the control mice. This result indicates that the motor coordination is impaired in *Dlic1^-/-^* mice.

**Figure3.**
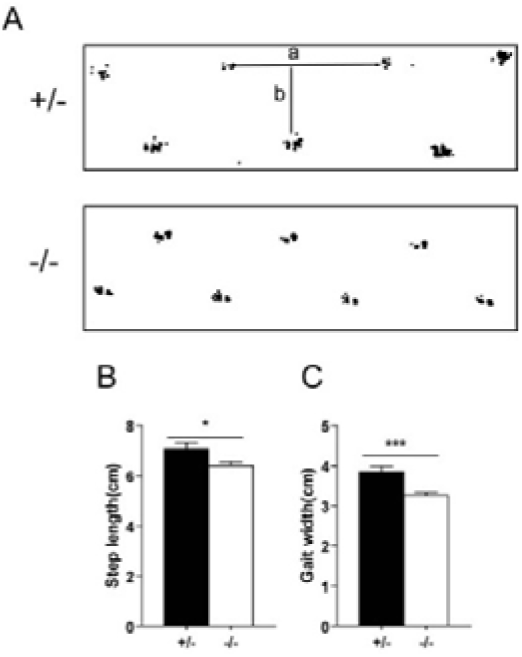
The *dlic1* deficiency affects mice footprint patterns. (A) The ink footprint illustrates the broad-based gait of the hind limbs of control and *dlic1* KO mice at 3 month. (B-C) Footprint patterns were quantitatively assessed for step length (B) and gait width (C). ***P < 0.001, *P < 0.05. Values represent the means ± SEM. n = 4 for each genotype.

### 3.3 Dlic1 deletion results in abnormal Purkinje cell dendrite arborization

In cerebellar development, Purkinje cells develop extensive dendritic arbors and synapse onto granule cells, controlling cell proliferation of the granule cell precursors[2]. The dendritogenesis of Purkinje cells is very important for the growth and pattern of cerebellar. To survey whether the defected cerebellar development in *Dlic1^-/-^* mice are related to the Purkinje cells (Figure4), we analyzed the morphology of the Purkinje cells with the antibody against calbindin, a specific marker for Purkinje cells. The result shows that the dendritogenesis of Purkinje cell was remarkably affected in *Dlic1^-/-^* cerebellum at P7 (Figure4B, C). The statistical analysis revealed that both the dendritic length and branch are severely reduced. Purkinje cells are the output cells in cerebellum and thus related to the motor dysfunction of *Dlic1^-/-^* mice.

**Figure4.**
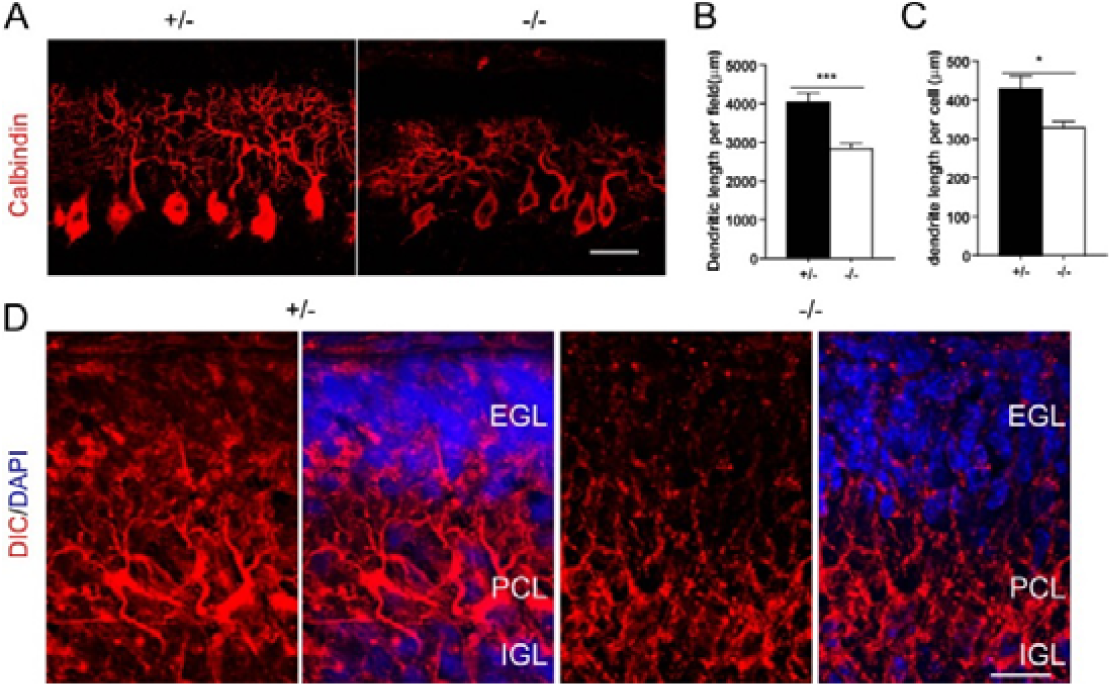
*Dlic1* deficiency impairs Purkinje cell dendritogenesis. (A) Representative images of P7 Purkinje cells in cryosections of cerebellum stained with anti-calbindin. (B-C) Quantitative analysis of total dendrite length per field (B) per cell (C) in P7 cerebellum slices. ***P < 0.001, *P < 0.05, ns, not significant. Values represent the means ± SEM. n = 3 for each genotype. (D) Decreased expression of DIC in *dlic1^-/-^* Purkinje dendrites from P4 mice. Scale bar: 20μm.

To examine the how the absence of *Dlic1* leads to the defected Purkinje development, we used IF to detect the dynein expression in *Dlic1^-/-^* cerebellum (Figure4D). The results show that the expression level of DIC is severely reduced in Purkinje dendrites in *Dlic1^-/-^* cerebellum. This strongly indicates that the dendritogenesis of Purkinje cells is impaired by the dynein reduction.

### 3.4 Dlic1 regulates neurite outgrowth in cerebellar granule cells

Formation of neuronal neurites is important for the postnatal development of cerebellum. After proliferation, granule cells extrude bipolar axons to form parallel fibers and move tangentially along the fibers in the inner granule layer[20]. To explore whether the granule cells contribute to the impaired *Dlic1^-/-^* cerebellar development, we examined the neurite growth of granule cells isolated from the cerebellar (Figure5). Cultured granule cells show progressive exogenous during the first several days preceding the development of shorter dendrites. In isolated granule neurons, the neurite length and the neurite branch number are reduced in *Dlic1^-/-^* granule cells(Figure5B). These results strongly suggest that dlic1 is involved in neurite extension in cerebellar granule cells.

**Figure5.**
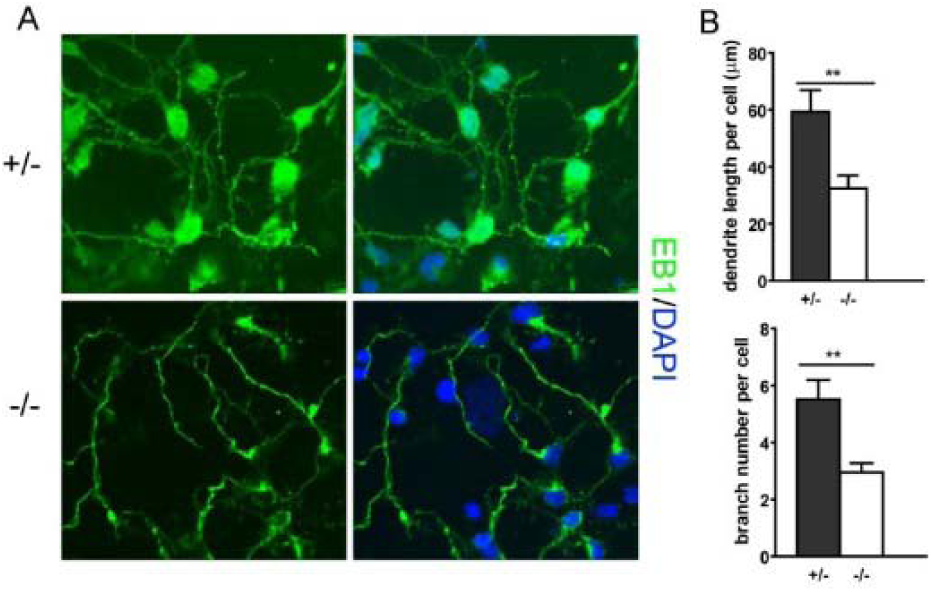
*Dlic1* deficiency impairs granule cell dendrite growth. (A)Immunostaining using EB1 antibody to mark the neurite of granule cells dissociated from P4 mice cerebellum and in vitro cultured for 36 hours. (B) Quantitative analysis of total dendrite length (up panel) and branch number (low panel) per cell. **P < 0.01, n = 3 mice for each genotype.

### 3.5 Dynein subunits were affected in Dlic1^*-/-*^ brain

In neurons, dynein complex is responsible for the retrograde transport from axon terminus to the cell body, which is required for neuronal survival and axonal plasticity[21].DLIC1 is a subunit of dynein complex helping DIC binding to DHC, LIS1 has been reported to be interact with DIC and help dynein complex binding to microtubules. To investigate how the deletion of *Dlic1* affected the dynein complex, we analyzed the assembling of dynein complex with the biochemical method.

First, we detected the protein level of DHC and DIC in whole brain lysate and found that the expression level of DHC and DIC were reduced when DLIC1 is deleted. The western blot analysis shows that the amount of LIS1 is not changed. The kinesin is also remained the same with control in the DLIC1 depleted brain.

Second, to investigate whether the interaction between DHC and DIC is affected by the deletion of DLIC1, we execute DIC-IP experiment, and observed an increased amount of the DHC and DLIC2 binding to DIC. The increased amount of DLIC2 binding on the DIC is probably because of the complementary effect: when *Dlic1* is depleted, some DHC and DIC are degraded, but the DLIC2 is not reduced. The DLIC2 binds to DHC and help the binding between DHC and DIC. Therefore, the interaction in the remaining DHC and DIC seems not impaired.

Third, to investigate whether the deletion of DLIC1 affected integrity of dynein complex in *Dlic1^-/-^* brain, we executed the sucrose gradient assay to analyze the dynein complex. In brief, whole cell Lysates of either WT or *Dlic1^-/-^* brain were separated by sucrose gradient centrifugation, the collected fractions were analyzed by immunoblotting (Fig. 6 A,B). The result shows that both the DIC and EB1 migrated to lower molecular weight fractions when *Dlic1* is deleted. This probably because the dynein complex could not link to the cargo when DLIC1 is deleted, the molecule weight of dynein complexes tends to be smaller [16]. This is consistent with the previous result that the interaction of dynein and cargos is decreased when *Dlic1* is deleted.

**Figure6.**
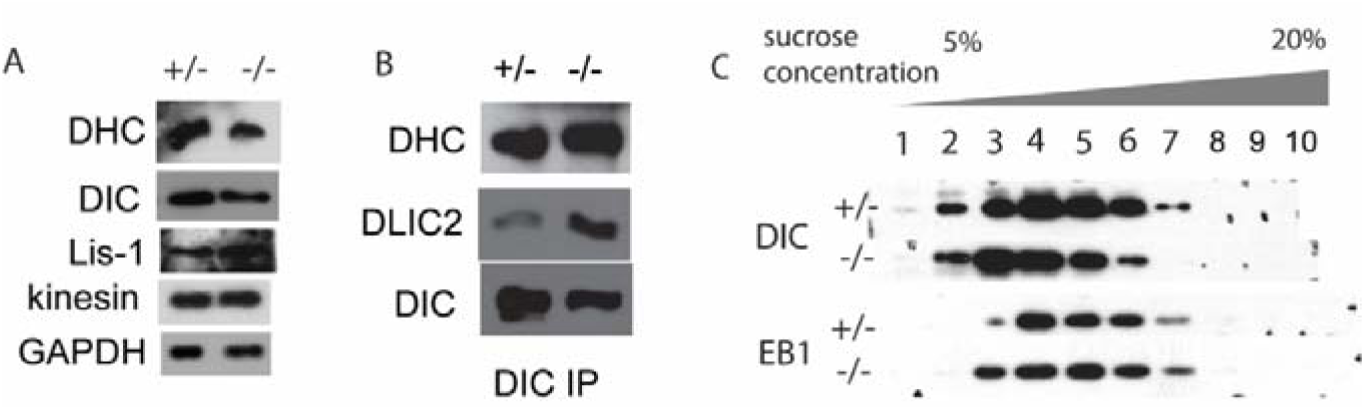
Biochemical analysis of dynein structure. (A) Western blot to detect some protein level in brain lysate. Anti-DHC, anti-DIC, Decreased expression of DHC and DIC in *dlic1^-/-^* Purkinje cell. Whole Brain were lysis and SDS-PAGe running and then western blotting with DHC, DIC in mouse brain. (B) Increased binding of DHC and DLIC2 to DIC when DLIC1 is depleted. Whole brain lysates was immunoprecipitated with anti-DIC then western blotting with anti-DHC and anti-DLIC2. (C) Sucrose gradient fraction analysis of dynein subunits. Whole cell lysate from mouse brain was loaded onto a 5-20% sucrose gradient and subject to ultra-centrifugation. The fractions were collected and analyzed by western blotting with anti-DIC and anti-EB1.

## 4. Discussion

This study addresses the function of *Dlic1* in cerebellar neurons both in vivo and in-vitro. Histological analysis demonstrated an evident size reduction and impaired foliation pattern in the *Dlic1^-/-^* cerebellum. Mice with depleted *Dlic1* display abnormal Purkinje cell morphogenesis and defective granule cell maturation. Consistent with the histological findings on cerebellar alterations, *Dlic1^-/-^* mice show reduced gait length and width. Therefore, our results demonstrate that *Dlic1* has a critical role in cerebellar development and function.

During the processes of cerebellar development, granule cells undergo profound morphological changes, including neurite initiation, neurite extension, and directed turning [22]. Neurite initiation and extension involves coordinated changes in actin and microtubule, which function to stabilize and maintain the neuritis [23].It is most likely that the affected neuron development in *Dlic1^-/-^* cerebellum is related with the defected dynein complex, which functions to. Further studies are required to elucidate how the redistributed signaling impaired the neuron development. The processes associated with neurite outgrowth are highly influenced by changes in the cellular environment.

Since *Dlic1* is required for cilium development [16], previous studies show that hedgehog pathway plays important role in cerebellar development[24]. Shh increased neurite outgrowth of cortical neurons grown on reactive astrocytes. Whether *Dlic1* deletion leads to cerebellar ataxia through the affected the cilium pathway is not known.

Previous studies have found that dynein light intermediate chain plays an important role in neuron development. Mice with a point mutation of *Dlic1*^*N235Y*^ display an increase in dendrite length in cortical neurons (with no changes in branching) and the number of dendrite branches in DRG neurons, in addition to increased anxiety-like behavior and altered gait [25]. However, *Dlic1^-/-^* mice exhibit reduced dendrite length and branch number in both Purkinje and granule cell. The differences arise because the *Dlic1*^*N235Y*^ mouse was thought to be a gain of function, and *Dlic1^-/-^* mouse we generated is a loss of function mutation. Our results were in accordance with the studies in Drosophila: mutating Drosophila *dlic1* gene results in reduced length and number of dendrite branches, and the mutation is also a loss of function [26], [27]. The previous study also shows that the deletion of dynein light intermediate chain impaired the interaction between DHC and DIC, but our results show that the deletion of dlic1 didn’t affect the interaction of remaining DHC and DIC. This is because there is another homolog *Dlic2* in the mouse.

Our previous data found that, in mouse CNS, *Dlic1* deletion leads to reduced total protein level of dynein, and the distribution of the remaining dynein complex in *Dlic1^-/-^* neurons is affected both in vivo and in vitro, indicating the universal role of *Dlic1* in dynein distribution.

In conclusion, we have clarified the important role of *Dlic1* in cerebellar development, elucidating the role of *Dlic1* will provide a novel tool for the future treatment of cerebellar ataxia.

## Acknowledgements

This work was supported by National Basic Research Program of China (2013CB945304, 2011CB510102, 2009CB526502, 2006CB806700); National Natural Science Foundation of China (31171406, 30971666, 30630043); Hi-tech Research and Development Program of China (2007AA022101); Key Projects Grant for Basic Research (08JC1400800) from the Science and Technology Committee of Shanghai Municipality; and the 211 and 985 projects of the Chinese Ministry of Education; This work was also supported by Excellent Ph.D. Student Research Grant of Fudan University to S. Kong. We thank Drs Chao Peng, Shunfei Yan, Xingrong Du for scientific discussion.

## Author contributions

SK, and WT. Conceived or designed the experiments; SK performed the experiments and analyzed the data; and SK wrote the manuscript.

